# Analysis of genome-wide differentiation between native and introduced populations of the cupped oysters *Crassostrea gigas* and *Crassostrea angulata*

**DOI:** 10.1101/292144

**Authors:** Pierre-Alexandre Gagnaire, Jean-Baptiste Lamy, Florence Cornette, Serge Heurtebise, Lionel Dégremont, Emilie Flahauw, Pierre Boudry, Nicolas Bierne, Sylvie Lapègue

## Abstract

The Pacific cupped oyster is genetically subdivided into two sister taxa, *Crassostrea gigas* and *C. angulata*, which are in contact in the north-western Pacific. The nature and origin of their genetic and taxonomic differentiation remains controversial due the lack of known reproductive barriers and morphologic similarity. In particular, whether ecological and/or intrinsic isolating mechanisms participate to species divergence remains unknown. The recent co-introduction of both taxa into Europe offers a unique opportunity to test how genetic differentiation maintains under new environmental and demographic conditions. We generated a pseudo-chromosome assembly of the Pacific oyster genome using a combination of BAC-end sequencing and scaffold anchoring to a new high-density linkage map. We characterized genome-wide differentiation between *C. angulata* and *C. gigas* in both their native and introduced ranges, and showed that gene flow between species has been facilitated by their recent co-introductions in Europe. Nevertheless, patterns of genomic divergence between species remain highly similar in Asia and Europe, suggesting that the environmental transition caused by the co-introduction of the two species did not affect the genomic architecture of their partial reproductive isolation. Increased genetic differentiation was preferentially found in regions of low recombination. Using historical demographic inference, we show that the heterogeneity of differentiation across the genome is well explained by a scenario whereby recent gene flow has eroded past differentiation at different rates across the genome after a period of geographical isolation. Our results thus support the view that low-recombining regions help in maintaining intrinsic genetic differences between the two species.

## Introduction

Broadcast-spawning marine invertebrates usually display high genetic similarity between adjacent populations, a consequence of combining high dispersal ability and large population sizes. Many such species, however, appear to be genetically subdivided into sibling species, cryptic species-pairs, ecotypes, or partially reproductively isolated populations (Knowlton, 1993, Bierne et al., 2011, Palumbi, 1994, Kelley et al., 2016, Hellberg, 2009, Gagnaire et al., 2015). The ability to score divergence at the genomic level in non-model organisms has recently revealed an increasing number of cases of cryptic species subdivision in broadcast-spawning marine invertebrates (e.g. Ravinet et al., 2015, Westram et al., 2014, Fraïsse et al., 2016, Rose et al., 2018). These recent findings confirm the view that semi-permeable barriers to gene flow between closely related taxa (Harrison & Larson, 2014, Harrison & Larson, 2016) are relatively frequent in the marine realm, and may explain short-distance genetic differentiation patterns that are seemingly in contradiction with species’ dispersal potential.

The existence of marine semi-isolated species pairs evolving in the “grey zone” of the speciation continuum (Roux et al., 2016), that is, before complete reproductive isolation, provides interesting opportunities to contribute to some highly debated questions in the field of speciation genomics. Speciation is a progressive process by which reproductive isolation barriers of various types progressively appear and combine their effects to reduce gene flow (Abbott et al., 2013). As long as speciation is not complete, diverging populations continue to evolve non-independently because some regions of their genome can still be exchanged. This results in a dynamic architecture of divergence characterized by temporal changes in the number, genomic distribution and magnitude of the genetic differences between incipient species. The speciation genomics approach investigates this architecture by characterizing heterogeneous genome divergence patterns, ultimately aiming at a detection of the loci directly involved in reproductive isolation (Feder et al., 2012). This strategy, however, has come up against a large diversity of evolutionary processes influencing the genomic landscape of species divergence (Ravinet et al., 2017, Wolf & Ellegren, 2017, Yeaman et al., 2016), some of which are not directly linked to speciation (Noor & Bennett, 2009, Nachman & Payseur, 2012, Cruickshank & Hahn, 2014, Burri, 2017). The field is now progressively moving toward a more mechanistic understanding of the evolutionary processes underlying heterogeneous genome divergence. However, the issue of distinguishing the impact of genetic barriers from the effect of confounding processes such as linked selection remains challenging (Burri, 2017). Because genetic divergence does not easily maintain in the face of gene flow in the absence of genetic barriers (Bierne et al., 2013), high gene flow species such as broadcast-spawning marine invertebrates offer valuable study systems for disentangling the mechanisms at play during speciation.

Here, we investigated the existence and the type of genetic barriers between divergent lineages of the Pacific cupped oyster, which has been taxonomically subdivided into two sister species, *Crassostrea gigas* and *C. angulata*. The two species are presumed to be parapatrically distributed in their native range in the north-western Pacific, but the location of putative contact zones remains largely unknown. Whether *C. gigas* and *C. angulata* truly represent biological species, semi-isolated species of populations of the same species also remains unclear. The two taxa can be cross-fertilized in the laboratory to form viable and fertile offspring (Huvet et al., 2001, Huvet et al., 2002, Takeo & Sakai, 1961), and some authors have proposed that they should be considered as a single species (Reece et al., 2008, López-Flores et al., 2004). On the other hand, the finding of genetic differences between *C. angulata* and *C. gigas* lead other authors to conclude that they form different but genetically closely related species (Boudry et al., 1998, Lapegue et al., 2004, Ren et al., 2010, Wang et al., 2014). One originality of the Pacific cupped oyster system is the co-introduction of both taxa into Europe. *C. angulata*, also called the Portuguese oyster, is presumed to have been non-voluntarily introduced by merchant ships during the 16^th^ century, probably from Taiwan (Boudry et al., 1998, Huvet et al., 2000) although the exact origins of introduced stocks remains unknown (Grade et al., 2016). Recent studies have shown that this species is widely distributed in Asian seas where it shows a high genetic diversity (Zhong et al., 2014b, Sekino & Yamashita, 2013, Wang et al., 2010, Hsiao et al., 2016), and also suggested a more complex history of transfers between Asia and Europe (Grade et al., 2016). The Pacific oyster, *C. gigas*, has been voluntarily introduced from Japan and British Columbia into Europe in the early 1970s, mainly to replace the Portuguese oyster in the French shellfish industry following a severe disease outbreak (Grizel & Héral, 1991). Since then, the two species are in contact in southern Europe and therefore have the potential to exchange genes in a new environment (Batista, et al. 2017; Huvet, et al. 2004). Gene flow in the context of marine invasions has mainly been studied between native and non-indigenous lineages (Saarman and Pogson 2015; Viard, et al. 2016) but rarely between two co-introduced genetic backgrounds in a new place. A notorious exception is the European green crab (*Carcinus maenas*) in the Northwest Atlantic (Darling, et al. 2008). However, although the genome-wide genetic differentiation has been studied in the introduced range (Jeffery, et al. 2017; Jeffery, et al. 2018), it has not been compared with the differentiation observed in the native range to date.

In the present study, we first generated new genomic resources in the Pacific oyster to improve the species genome assembly and characterize chromosomal variation in recombination rate. Then, we tested the existence of genetic barriers between *C. angulata* and *C. gigas* by searching for genomic regions that remain differentiated in the presence of gene flow, accounting for the demographic divergence history of the species. We also evaluated whether ecological divergence driven by local adaptations is the main factor maintaining species divergence in the native range. We hypothesized that if this is the case, the different ecological conditions encountered by the two species in Europe would have reshaped the original genomic landscape of species divergence existing in the native Asian range. Finally, we attempted to relate genome-wide divergence patterns to underlying evolutionary processes including demography, selection and genomic constraints.

## Material and Methods

### Biological material for the mapping populations

Two F2 families (second generation of biparental crosses) were used for genetic map reconstruction in *C. gigas*. These families were obtained by experimental breeding as part of the MOREST project (Boudry, et al. 2008; Dégremont, et al. 2010) by crossing families selected for resistance (*R*) or susceptibility (*S*) to summer mortality, which were subsequently found to have respectively higher and lower resistance to the herpesvirus OsHV-1 (Dégremont 2011). The F2-19 family was generated through biparental crossing between an *S* female and an *R* male to produce a F1 family from which a single sib-pair was randomly sampled to produce the F2 progeny. The F2-21 family was obtained under the same mating scheme but starting with an *R* female and a *S* male. A total of 293 and 282 F2 progenies were used for map reconstruction in family F2-19 and F2-21, respectively. For each individual, whole genomic DNA was isolated from muscle tissue using the QIAamp DNA minikit (Qiagen). DNA was checked for quality by electrophoresis on agarose gel and then quantified using the Quant-iT PicoGreen dsDNA assay kit (Life Technologies).

### Genotyping panel for low-density map construction

We genotyped F1 parents and their F2 progenies in both families (F2-19: 2 F1 and 293 F2; F2-21: 2 F1 and 282 F2) using a panel of 384 SNPs previously developed in *C. gigas* (Lapègue et al., 2014). SNP genotyping was performed using the Golden Gate assay and analyzed with the Genome Studio software (Illumina Inc). In addition, 42 microsatellite markers were also genotyped according to published protocols (Li et al., 2003, Taris et al., 2005, Yamtich et al., 2005, Li et al., 2010, Sauvage et al., 2010) in order to include markers from previous generation linkage maps in *C. gigas* (Hubert & Hedgecock, 2004, Sauvage et al., 2010).

### RAD genotyping for high-density map construction

We selected 106 progenies from each family as well as their four F1 parents for RAD library construction following the original protocol (Baird et al., 2008). Briefly, 1 μg of genomic DNA from each individual was digested with the restriction enzyme *Sbf*I-HF (New England Biolabs), and then ligated to a P1 adapter labeled with a unique barcode. We used 16 barcodes of 5-bp and 16 barcodes of 6-bp long in our P1 adapters to build 32-plex libraries. Seven pools of 32 individuals were made by mixing individual DNA in equimolar proportions. Each pool was then sheared to a 350 pb average size using a Covaris S220 sonicator (KBiosciences), and size-selected on agarose gel to keep DNA fragments within the size range 300-700 pb. Each library was then submitted to end-repair, A-tailing and ligation to P2 adapter before PCR amplification for 18 cycles. Amplification products from six PCR replicates were pooled for each library, gel-purified after size selection and quantified on a 2100 Bioanalyzer using the High Sensitivity DNA kit (Agilent). Each library was sequenced on a separate lane of an Illumina HiSeq 2000 instrument by IntegraGen Inc. (France, Evry), using 100-bp single reads.

We used the program *Stacks* (Catchen et al., 2013, Catchen et al., 2011) to build loci from short-read sequences and determine individual genotypes. Raw sequence reads were quality filtered and demultiplexed using *process_radtags.pl* before being trimmed to 95 bp. We explored different combinations of parameter values for the minimum stack depth (-m) and the maximum mismatch distance (-M) allowed between two stacks to be merged into a locus. We found that the combination - m 3 -M 7 represented the best compromise to avoid overmerging loci, while providing an average number of 2.3 SNPs per polymorphic RAD locus (Supplementary Figure S1) which is consistent with the high polymorphism rate in *C. gigas* (Sauvage et al., 2007, Zhang et al., 2012). Individual *de novo* stack formation was done with *ustacks* (-m 3 -M 7 -r -d). We then built a catalog of loci using all individuals from both families with *cstacks* (-n 7) and matched back all the samples against this catalog using *sstacks*. After this step, the two families were treated as two separate populations to produce a table of observed haplotypes at each locus in each family using *populations*.

We developed a Bayesian approach that uses information from progenies’ genotypes to correct for missing and miscalled genotypes in parental samples. The probability of a given combination of parental genotypes (*G*_*P*_) conditional on the genotypes observed in their descendants (*G*_*D*_) is given by

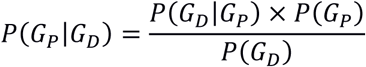

where the probability of the observed counts of progenies’ genotypes (*N*_AA_, *N*_AB_, *N*_AC_, *N*_AD_, *N*_BB_, *N*_BC_, *N*_BD_) conditional on the genotypes of their parents is drawn from a multinomial distribution that takes different parameter values for each of the six alternative models of parental genotypes (i.e. crosses AA×AA, AA×AB, AB×AB, AA×BB, AB×AC and AB×CD). Each RAD locus was treated independently using observed haplotypes to determine the best model of parental haplotype combination. When the actual haplotypes called in parents did not match the best model, a correction was applied to restore the most likely combination of parental haplotypes, taking read depth information into account. Data were finally exported in *JoinMap 4* format with population type CP (van Ooijen, 2006). The approach for correcting missing and miscalled haplotypes in parental samples was coded into R.

### Genetic map construction

A new Pacific oyster linkage map was constructed in four successive steps: (*i*) First, a low-density map was built for each family using the SNP and microsatellite markers dataset. These two maps were used to provide accurate ordering of markers, since both mapping populations comprised approximately 300 individuals with very few missing genotypes (0.5%). (*ii*) In a second step, we tried to reach a consensus order for the markers that were included in the two previous maps, in order to determine a set of anchor loci for each linkage group. (*iii*) Third, we included RAD markers and determined the order of all loci in each mapping family after setting a fixed order for the anchor loci. (*iv*) Finally, we integrated the two high-density linkage maps to produce a consensus genetic map for the Pacific oyster.

All linkage mapping analysis were performed using *JoinMap 4* (van Ooijen, 2006). Markers showing significant segregation ratio distortion after Bonferroni correction (p < 0.05) were removed from the analysis. Markers were grouped using an independence LOD threshold of 16 for the SNP and microsatellite marker datasets and a LOD threshold of 10 for the RAD marker datasets. Additional ungrouped markers were assigned at a LOD threshold of 5 using their strongest cross-link information. We used the regression mapping algorithm to build the maps using a recombination frequency threshold of 0.4, a minimal LOD score of 1 and a goodness-of-fit jump value of 5. The ordering of markers was optimized using the ripple function after each added locus. Map distances in centiMorgans (cM) were calculated from recombination frequencies using Kosambi’s mapping function.

### Identification of chimeric scaffolds and reassembly using BAC-end scaffolding

In order to detect chimeric scaffolds in the oyster_v9 assembly (Zhang et al., 2012), the consensus sequences of the markers included in the new linkage map were blasted against the reference genome. An E-value threshold of 10^−30^ and a minimum identity threshold of 90% were set to retain only highly significant matches. Assembly errors were identified by scaffolds anchored to more than a single linkage group. Chimeric scaffolds were subsequently splitted at all stretches of Ns connecting adjacent contigs.

In order to improve the scaffolding of the Pacific oyster genome, we sequenced 73,728 BAC clones of 150 kb average insert size (Gaffney 2008) from both ends using Sanger sequencing at the Genoscope facility (Evry, France). This resulted in 60,912 cleaned full-length (i.e. >1300 bp) BAC-End sequences (BES) pairs. Each sequence from each pair was trimmed to only conserve 1000 bp between positions 10 and 1010, and clipped into 19 evenly-spaced 100-mers overlapping by 50 bp. This clipping procedure aimed at constructing a high quality short-read paired-end dataset from our BES dataset.

Clipped BES were used for an additional round of contig extension and scaffolding with *SSPACE* (Boetzer et al., 2011). For each LG, we used unambiguously anchored scaffolds and contigs originating from splitted scaffolds matching to this LG. Clipped BES were aligned to scaffolds using *Bowtie2* (Langmead & Salzberg, 2012), allowing at most 3 mismatches per 100-bp read. Scaffolding parameters to *SSPACE* were set to a minimum of 5 links (-k) to validate a new scaffold pair and a maximum link ratio of 0.7 (-a). Scaffolding was only permitted between scaffolds of at least 1000 bp, using an allowed insert size range of 75-225 kb.

### Scaffold anchoring to the linkage map

We searched for non-ambiguous associations between the new set of extended scaffolds and markers included in the consensus linkage map using the same blasting procedure as prior to genome reassembly. We constructed pseudo-chromosomes by positioning scaffolds along each linkage group using *Harry Plotter* (Moretto et al., 2010). Because the genetic resolution of our consensus map was still limited by the size of our mapping populations (about 100 individuals each), many scaffolds were anchored by a single marker or had too few markers to determine their orientation. Therefore, we did not determine scaffold orientation. Scaffolds that were anchored to multiple markers were positioned using the average cM value of their anchor loci. Unmapped scaffolds were placed in an artificial chromosome named UN.

### Local recombination rate estimation

We used the R package *MareyMap v1.2* (Rezvoy et al., 2007) to estimate local recombination rates along the genome, by comparing the consensus linkage map and the physical map for each pseudo-chromosome. The relationship between genetic and physical distances was first visualized to remove outlier markers resulting from scaffold misplacement within pseudo-chromosomes. We then used the Loess interpolation method which estimates recombination rates by locally adjusting a 2^nd^ degree polynomial curve, setting the span parameter value to 0.25.

### RAD genotyping of natural populations

We sampled four wild populations of *C. gigas* and *C. angulata* from both their native Asian and introduced Atlantic areas. We used 24 individuals of *C. gigas* from both Japan (native) and French Britany (introduced), and 24 individuals of *C. angulata* from both Taiwan (native) and Portugal (introduced). We prepared and sequenced four 24-plex RAD libraries using the same protocol as described above.

Cleaned demultiplexed reads were mapped to each newly assembled pseudo-chromosome using *Bowtie2* (Langmead & Salzberg, 2012) with the very-sensitive option, allowing a maximum of 7 mismatches per alignment. SNPs were called from aligned reads with *Stacks* using a minimum read depth of 5x per individual per allele (Catchen et al., 2011, Catchen et al., 2013). The correction module *rxstacks* was used to re-evaluate individual genotypes and exclude low-quality variants with a cutoff log-likelihood value of −500. Only RAD loci that were successfully genotyped in at least 80% of the samples in each population were retained for subsequent population genomic analyses.

### Population genomic analyses

We used *VCFtools* v0.1.11 (Danecek et al., 2011) to apply within-population filters to exclude loci showing more than 4 missing genotypes over 24 individuals, as well as markers showing departure to Hardy-Weinberg equilibrium within at least one population using a p-value cutoff of 0.01. Nucleotide diversity (π), computed as the average number of pairwise differences, was estimated from retained loci for each population within 150 kb windows. Genetic differentiation between all possible pairs of populations was estimated SNP by SNP as well as in 150 kb windows using *F*_ST_ (Weir & Cockerham, 1984). Within- and between-population components of genetic diversity were decomposed using a discriminant analysis of principal components (dAPC), in order to maximize genetic variation between populations while minimizing within-population variance. The dAPC analysis performed with the R package *Adegenet* (Jombart, 2008), using the global SNP dataset containing the two populations from both species with only one randomly selected SNP per RAD locus.

In order to evaluate the influence of recombination rate variation on genetic differentiation between species, we used a nonparametric quantile regression approach. Recombination rate and *F*_ST_ values (averaged between the Asian and European species pair) were averaged in 500 kb windows to increase the number of informative sites per window. We excluded windows with recombination rate values exceeding 9 cM/Mb (i.e. corresponding to the 98^th^ percentile of the distribution of estimated recombination rate). This cutoff aimed at removing outlying recombination rate values estimated along chromosome IX (Figure 1). The 97.5^th^ quantile regression fit of the distribution of *F*_ST_ as a function of recombination rate was computed at each of 10 equally spaced recombination intervals distributed over the range of recombination values.

**Fig. 1.**
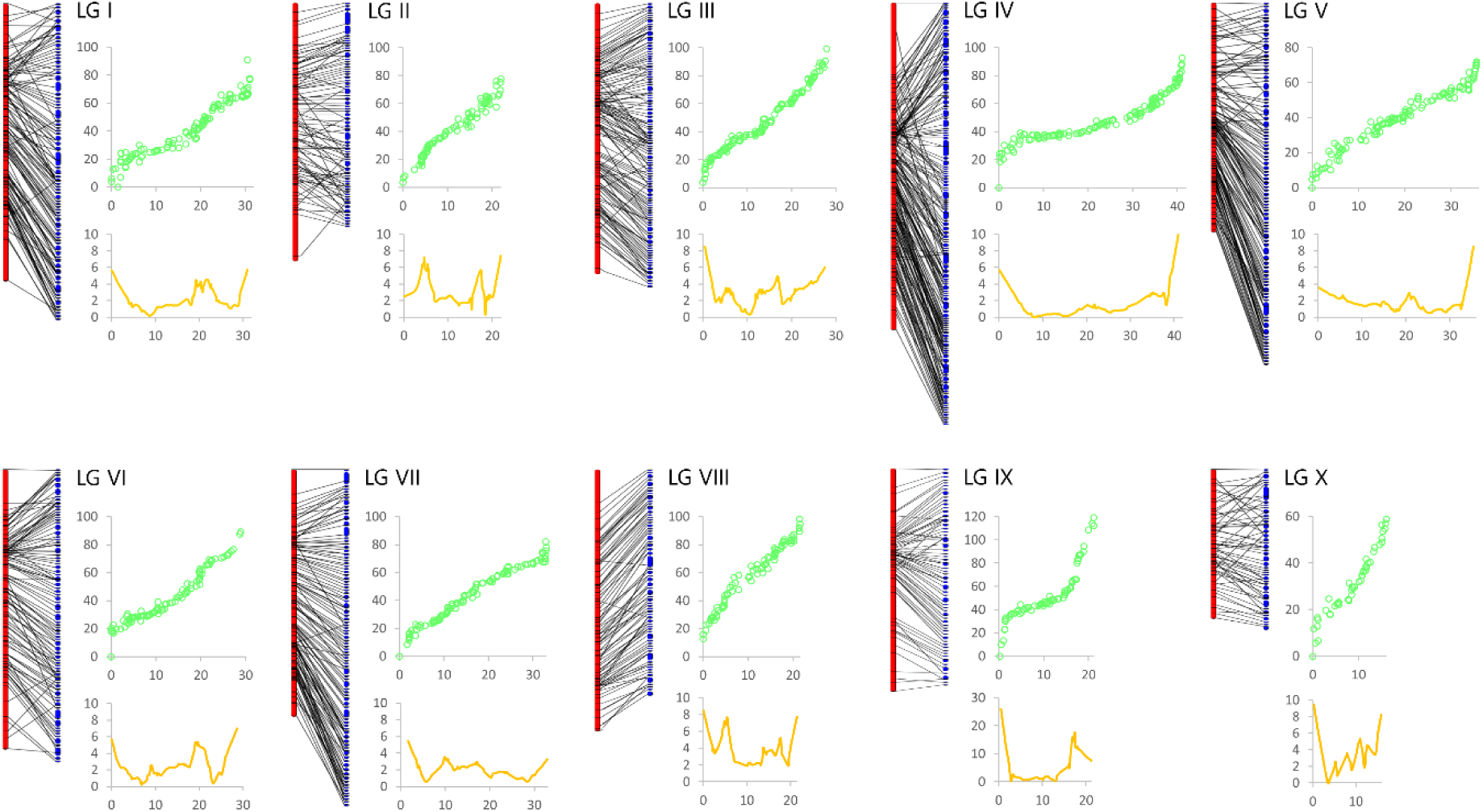
Anchoring of newly edited scaffolds (blue) into pseudo-chromosomes using the markers included in the 10 linkage groups (red) of the new consensus linkage map. Comparison of the physical (x axis, Mb) and the genetic (y axis, cM) maps are provided for each linkage group (green points, outlier values removed), as well as the local recombination rate (orange line) estimated using the Loess interpolation method in *MareyMap*.

### Inference of the demographic divergence history

We inferred the demographic divergence history between *C. angulata* and *C. gigas* in both their native and introduced ranges using a version of the program *δaδi* (Gutenkunst et al., 2009) modified by (Tine et al., 2014). The observed joint allele frequency spectrum (JAFS) of each population pair was obtained by randomly selecting one SNP for each pair of RAD loci associated to the same restriction site in order to avoid the effect of linkage between SNPs, and then by projecting the data to 20 chromosomes per population to reduce the JAFS size. We first considered four classical models of divergence including models of strict isolation (SI), isolation-with-migration (IM), ancient migration (AM) and secondary contact (SC). We also considered extensions of the three divergence-with-gene-flow models assuming two categories of loci occurring in proportions *P* and 1-*P* and having different migration rates: IM2*m*, AM2*m* and SC2*m* (Tine et al., 2014). These models offer simplified but more realistic representations of the speciation process, by enabling loci to be either neutrally exchanged between populations, or to have a reduced effective migration rate to account for the direct and indirect effects of selection. Each model was fitted to the observed JAFS (singletons were masked) using three successive optimization steps: hot simulated annealing, cold simulated annealing and BFGS. Comparison among models was made using the Akaike Information Criterion (AIC).

## Results

### Construction of an integrated high-density linkage map

The microsatellite and SNP datasets used for constructing the low-density linkage map contained 133 informative markers in family F2-19 (293 progeny, Table S1) and 137 informative markers in family F2-21 (282 progeny, Table S2). Ten linkage groups were found in each family in agreement with the haploid number of chromosomes of the species, and the two maps generated by the first round of regression mapping respectively contained 98 and 105 mapped markers in family F19 and F21. The integrated map obtained after identifying homologous LG pairs between families contained 136 markers successfully ordered after the first round of regression mapping. This sex-averaged consensus low-density linkage map displayed a strong collinearity with the two maps from which it was derived (Supplementary Figure S2, left panel), indicating that the ordering of markers was highly consistent between the two families. We thus used it as an anchoring map to support the construction of the high-density RAD-linkage map.

The average number of filtered RAD-Sequencing reads obtained per individual was similar between family F2-19 (1.8×10^6^) and F2-21 (1.9×10^6^). After correcting for missing and miscalled haplotypes in parental samples, the number of informative markers was 1278 in family F2-19 and 996 in family F2-21 (Table S3, Table S4). A total of 1855 markers were assigned to linkage groups (1231 in family F19 and 935 in family F21, 311 in common), and the number of markers per linkage groups was highly positively correlated between families (R²=0.71, Supplementary Figure S3). Using the anchoring map to set fixed orders, the two RAD maps generated after three rounds of regression mapping respectively contained 1023 and 705 mapped markers in family F19 and F21, respectively. The final combination of these two maps resulted in a sex-averaged consensus map containing 1516 markers (Table 1, Table S5, and Supplementary Figure S2, right panel). The total map length was 965 cM and the average spacing between two neighboring markers was 0.64 cM. We compared this new linkage map to the Pacific oyster’s second generation linkage map (Table S7 from Hedgecock et al., 2015) using scaffolds from the oyster reference genome (oyster_v9, Zhang et al., 2012) as intermediate. Using 278 pairs of markers colocalized to the same scaffolds, we found a good collinearity between the two maps, as illustrated by linkage group-wise correlations between recombination distances (0.35 < R² < 0.90, Supplementary Figure S4).

**Table 1.** Summary statistics for the 10 linkage groups of the new C. gigas high-density linkage map and the reassembled physical genome.

| LG | Correspondance with Hedgecock et al. 2015 | Length (cM) | Nb of markers F2-19 | Nb of markers F2-21 | Total nb of markers | marker spacing | physical length of anchored scaffolds | recombination rate (cM/Mb) | recombination rate MareyMap (cM/Mb) |
| --- | --- | --- | --- | --- | --- | --- | --- | --- | --- |
| I | 1 | 90.92 | 86 | 88 | 158 | 0.58 | 31113268 | 2.92 | 2.61 |
| II | 4 | 94.23 | 89 | 56 | 117 | 0.81 | 21949604 | 4.29 | 3.65 |
| III | 5 | 99.17 | 166 | 69 | 229 | 0.43 | 27979720 | 3.54 | 3.40 |
| IV | 7 | 116.98 | 149 | 103 | 226 | 0.52 | 41226161 | 2.84 | 2.29 |
| V | 10 | 72.01 | 95 | 74 | 163 | 0.44 | 35636424 | 2.02 | 2.18 |
| VI | 3 | 98.87 | 143 | 60 | 132 | 0.75 | 28856184 | 3.43 | 2.71 |
| VII | 8 & 2 | 88.78 | 113 | 71 | 157 | 0.57 | 33248857 | 2.67 | 2.36 |
| VIII | 9 | 108.20 | 64 | 68 | 131 | 0.83 | 22447078 | 4.82 | 4.13 |
| IX | 6 | 136.51 | 94 | 58 | 124 | 1.10 | 21192597 | 6.44 | 5.47 |
| X | 6 | 58.89 | 24 | 58 | 79 | 0.75 | 16594032 | 3.55 | 3.65 |
| ALL | - | 964.55 | 1023 | 705 | 1516 | 0.64 | 280243925 | 3.44 | 3.05 |

### Pseudo-chromosome assembly

A majority (78.3%) of the markers included in the new consensus RAD linkage map matched to the oyster reference genome (oyster_v9, Zhang et al., 2012). Among the 592 scaffolds that were matched with mapped markers, 327 (55.2%) contained a single mapped marker, 127 (21.5%) contained two markers, and 138 (23.3%) contained three to eleven markers (Table S6). We found 117 (44.2%) chimeric scaffolds mapping to different linkage groups among the 265 scaffolds containing at least two markers. Splitting chimeric scaffolds into smaller contigs at poly-N stretches decreased the assembly N50 size from 401 kb to 218 kb.

Contig extension and scaffolding using clipped BES allowed the merging of 4162 contigs. The mean insert size of mapped BES was 116 kb, which was consistent with the mean insert size of the BAC end library. The resulting N50 of new scaffolds increased by 47% to 320 kb (Table S7). A total of 773 scaffolds from this new set of extended scaffolds were anchored to 1161 loci of the consensus linkage map. Among them, 531 (68.7%) contained a single mapped marker, 153 (19.8%) contained two markers, and 89 (11.5%) contained three to seven markers (Table S8). The pseudo-chromosomes assembly obtained had a length of 280.2 Mb (Figure 1), representing 50.1% of the total length of the oyster_v9 assembly (Zhang et al., 2012).

### Local recombination rate

The comparison between the pseudo-chromosome assembly and the consensus linkage map revealed variation in local recombination rate along chromosomes, with generally increased values toward linkage group extremities compared to central chromosomal regions (Figure 1). The average genome-wide recombination rate estimated with *MareyMap* was 3.05 cM/Mb, a value close to the total map length divided by the size of the assembly (3.44 cM/Mb). The real value, however, may be twice lower considering that the pseudo-chromosome assembly only represents 50% of the total genome size. The average chromosome-wide recombination rate (Table 1) was negatively correlated with chromosome length (R² = 0.61, Figure 2), as expected under strong chiasma interference.

**Fig. 2.**
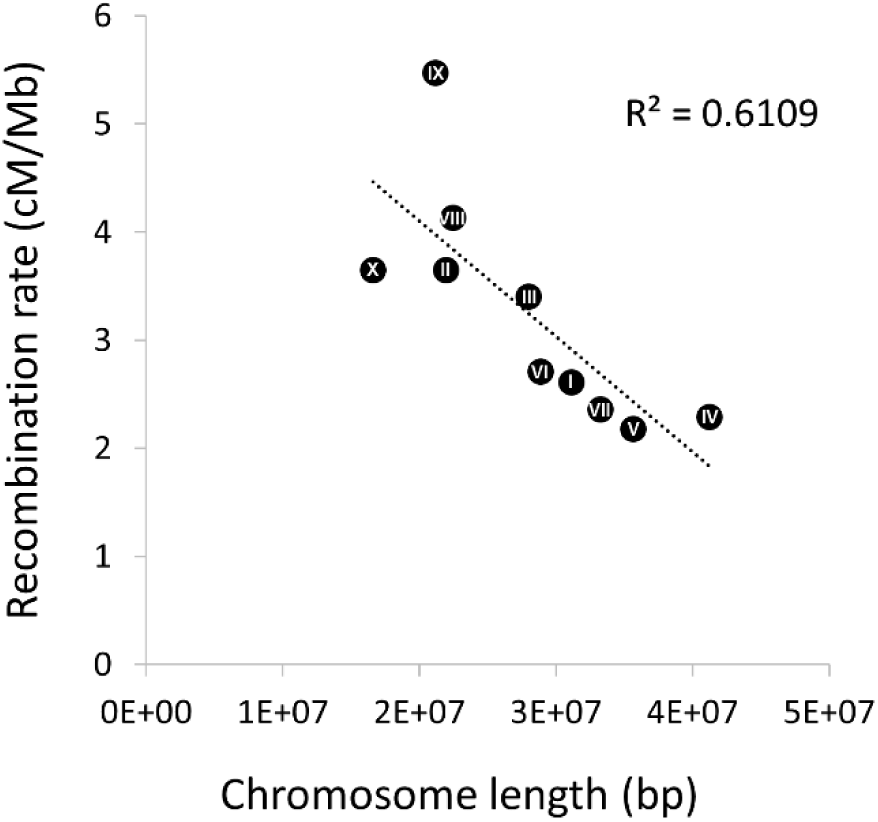
Negative correlation between the length of newly assembled pseudo-chromosomes and the average chromosome-wide recombination rate assessed with MareyMap. Chromosome lengths and raw estimates of recombination rate are based on the physical chromosome lengths effectively covered by scaffolds. Rescaled values taking into account the presence of unmapped scaffolds can be obtained by multiplying (dividing) chromosome length (recombination rate) values by two (see text).

### Genetic diversity and differentiation

The average number of sequence reads retained per individual for calling genotypes was 3.1×10^6^. After filtering for missing genotypes, HWE and a minor allele frequency of 1%, we kept a total of 10,144 SNPs from 1,325 RAD loci for downstream genetic diversity analyses. Within-population nucleotide diversity was elevated in both species and showed very similar levels between native and introduced populations (*C. angulata*: *π*_KEE_ = 0.0095, *π*_SAD_ = 0.0096; *C. gigas*: *π*_JAP_ = 0.0099, *π*_LAF_ = 0.0101). The genome-wide averaged differentiation assessed by *F*_ST_ between native and introduced populations from the same species was higher in *C. angulata* (*F*_ST KEE-SAD_ = 0.0179) compared to *C. gigas* (*F*_ST JAP/LAF_ = 0.0130). The mean genetic differentiation between species was very similar between Asia (*F*_ST KEE/JAP_ = 0.0440) and Europe (*F*_ST SAD/LAF_ = 0.0459).

The between-population genetic structure revealed by the dAPC (Figure 3A) separated the two species along the first PC axis (explaining 15.78% of total variance) and the Asian and European populations of each species along the second axis (2.54% of total variance). The European populations of *C. angulata*, and to a lesser extent *C. gigas*, were slightly shifted toward the center of the first PC axis, indicating an increased genetic similarity of the two species in Europe.

**Fig. 3.**
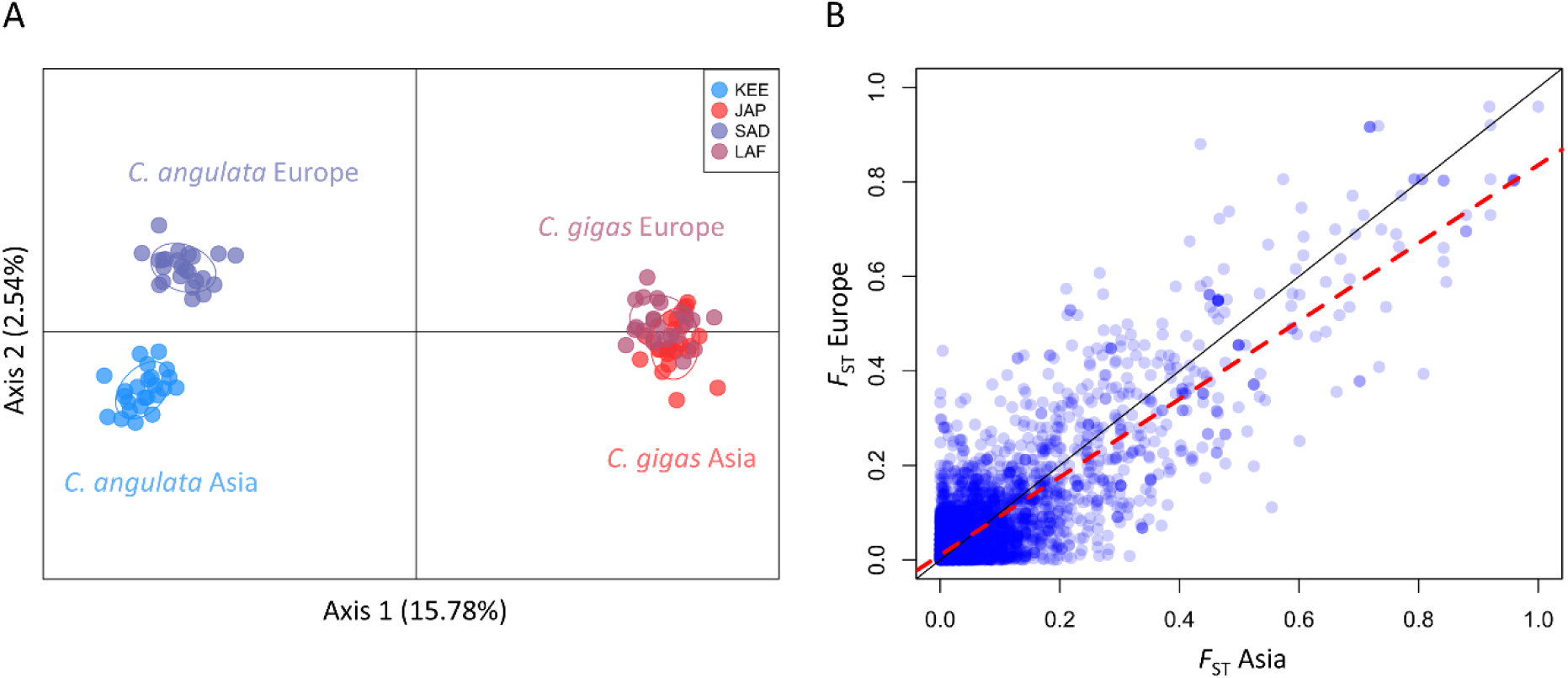
**(A)** dAPC plot of the four populations of Pacific cupped oysters assessed with 1,325 SNPs (one SNP per RAD), with C. angulata from Asia (KEE, light blue) and Europe (SAD, dark blue), and C. gigas from Asia (JAP, light red) and Europe (LAF, dark red). **(B)** Genetic parallelism in the level of differentiation between species in Asia (x axis) and Europe (y axis) measured at 10,144 SNPs. The black line shows the y = x equation line and the red dashed line corresponds to the linear regression (F_ST Europe_ = 0.83 × F_ST Asia_ + 0.01, R^2^ = 0.71, P > 10^−10^).

Genetic differentiation between species was highly heterogeneous across the genome, with regions of elevated differentiation alternating with genetically undifferentiated regions (Figure 4). The maximum *F*_ST_ value between *C. gigas* and *C. angulata* was 1 in Asia and 0.96 in Europe. Between-species genetic differentiation at individual SNPs was highly positively correlated between Asia and Europe, although it was on average lower in Europe (Figure 3B).

**Fig. 4.**
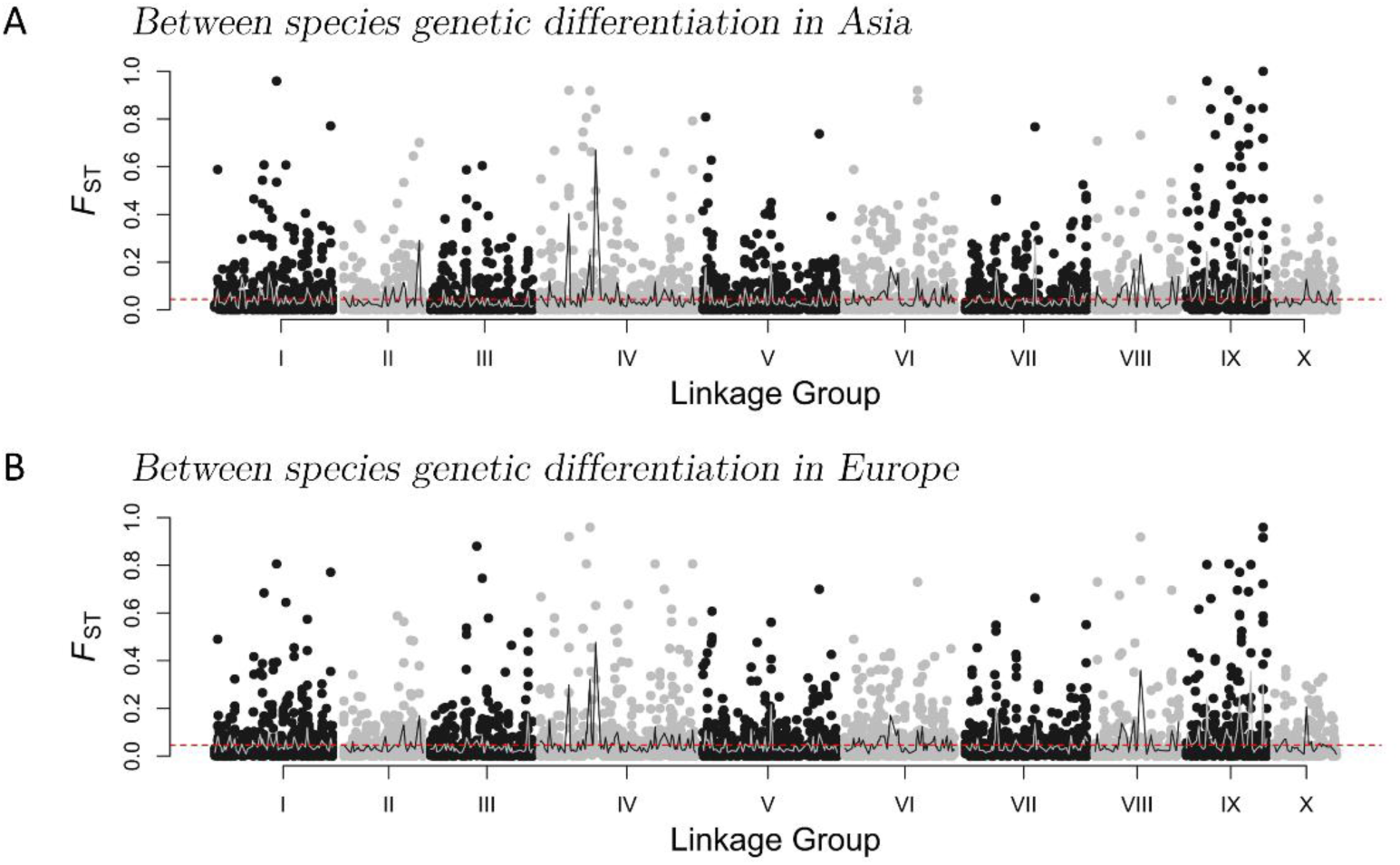
Genomic landscape of between-species genetic differentiation (F_ST_) measured at 10,144 SNPs (alternatively represented by black and grey points for odd and even chromosome numbers for convenience) along the ten Pacific oyster linkage groups. The red dotted line shows the genome-wide average F_ST_ and the grey line shows the local average F_ST_ calculated in 150 kb windows. Genetic differentiation between C. angulata and C. gigas is showed **(A)** in Asia and **(B)** in Europe.

Chromosomal variation in average genetic differentiation between species was related to the local recombination rate. More precisely, we found a decreasing trend in the maximal amount of differentiation (measured by the 97.5^th^ quantile of the distribution) with increasing local recombination rate (Figure 5). Moreover, within-species nucleotide diversity was significantly positively correlated with the local recombination rate (Supplementary Figure S5).

**Fig. 5.**
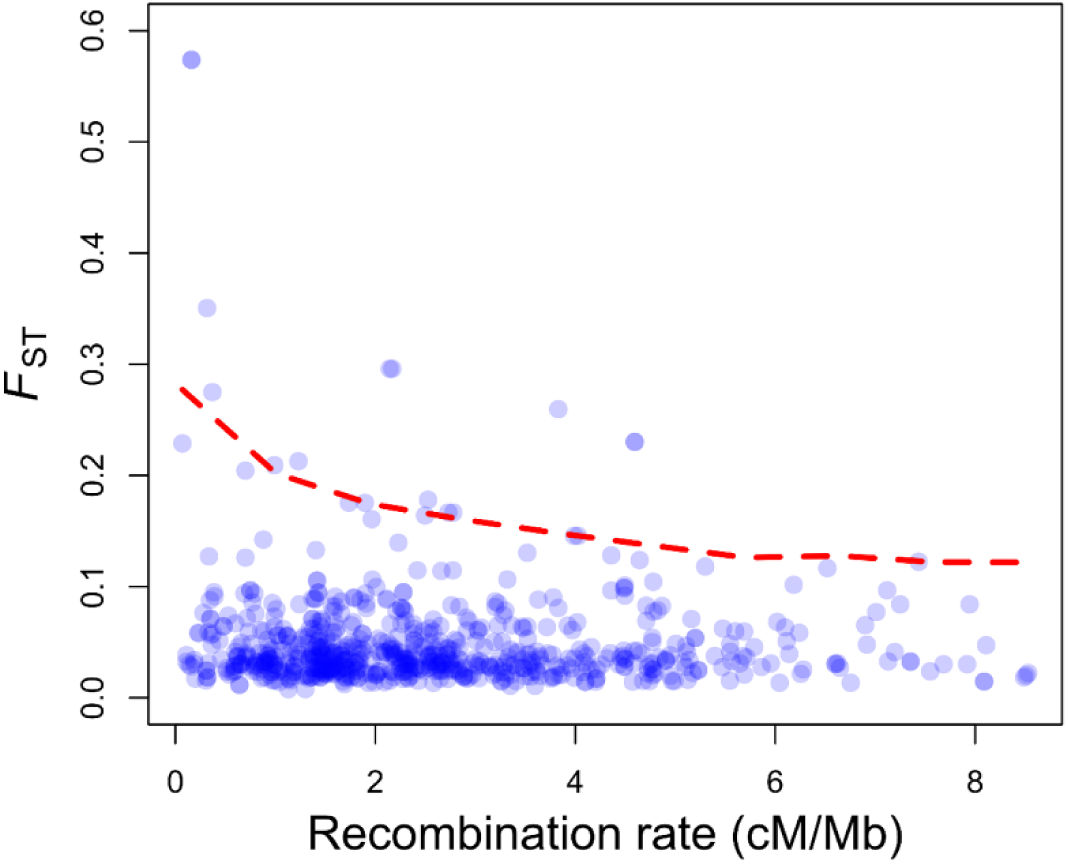
Distribution of between-species genetic differentiation (*F*_*ST*_) as a function of the local recombination rate assessed with *MareyMap*. Each point represents an estimate of *F*_*ST*_ and recombination rate averaged in a 500 kb window. Outlier recombination rate values exceeding 9 cM/Mb were excluded. The red dashed line represents the nonparametric 97.5^th^ quantile regression fit of *F*_*ST*_ as a function of the local recombination rate.

### Divergence history

The demographic history of divergence between *C. angulata* and *C. gigas* inferred with *δaδi* showed evidence for a secondary contact in both Asia and Europe (Table 2). The JAFS presented in Figure 6 show a high proportion of shared polymorphisms at high frequency (in the upper right and lower left corners) which is a characteristic footprint of secondary introgression that is not expected under a strict isolation model (Alcala et al., 2016, Roux et al., 2016) and explain the good support for the SC model. Moreover, varying rates of introgression among loci (*i.e.* the SC2*m* model) was strongly supported for both native and introduced species pairs. This model received a good statistical support compared to the six other alternative models (Akaike weights > 0.95), and its goodness-of-fit showed almost no trend in the distribution of residuals for both species pairs (Figure 6).

**Table 2.**
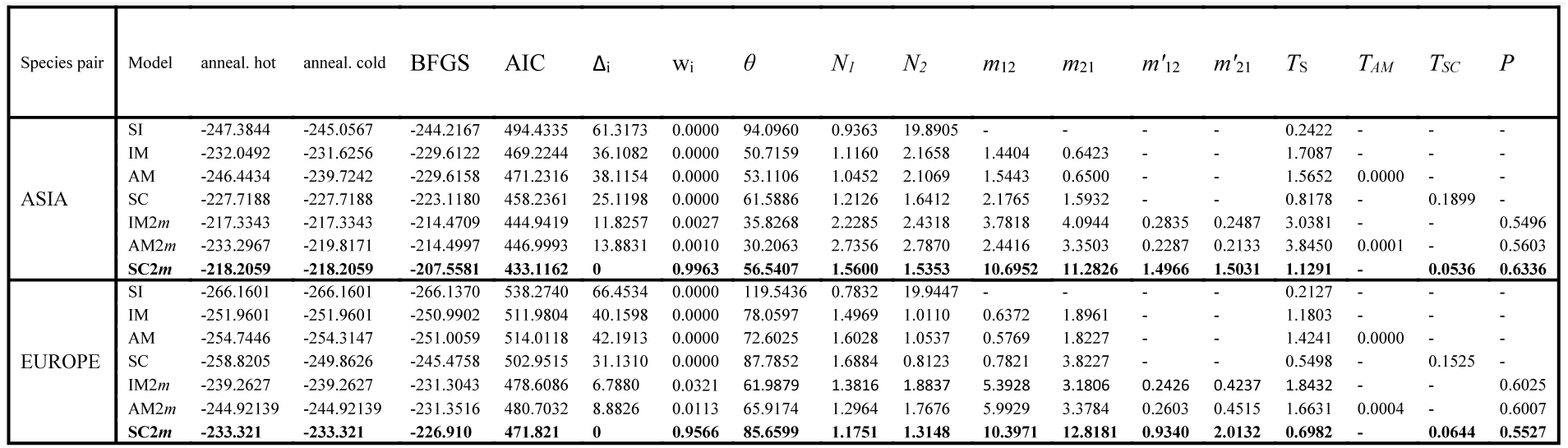
Results of model fitting for seven alternative models of divergence between C. angulata and *C. giga* likeliho anneali model (Follow *angulat* genom migrati within **2.** Results of model fitting for seven alternative models of divergence between *C. angulata* and *s* in Asia and Europe. In the order of appearance in the table: the model fitted, the maximum likelihood estimate over 20 independent runs after 3 successive optimisation steps: simulated annealing hot, cold and BFGS, the Akaike Information Criterion, the difference in AIC with the best SC2*m*), Akaike weight, and *theta* parameter for the ancestral population before split. Following are the inferred values for the model parameters (scaled by *theta*): the effective size of *C. angulata* (*N*_1_) and *C. gigas* (*N*_2_) populations, the migration rates for the two categories of loci in the genome (category 1: *m*_12_ and *m*_21_, category 2: *m’*_12_ and *m’*_21_), the duration of the split (*T*_S_), of ancestral migration (*T*_AM_) and secondary contact (*T*_SC_) episodes, and the proportion (*P*) of the genome falling category 1 (experiencing migration rates *m*_12_ and *m*_21_).

**Fig. 6.**
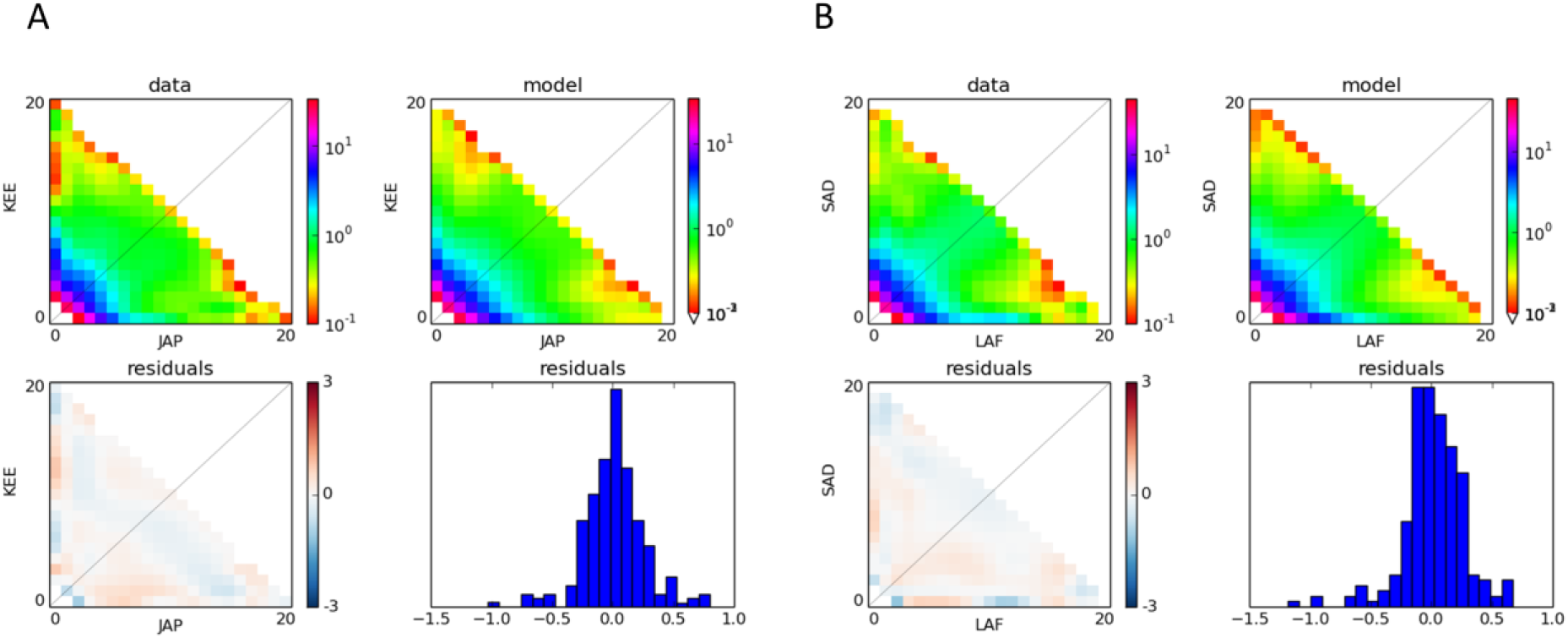
Demographic divergence history of *C. angulata* and *C. gigas* in Asia (**A**) and Europe (**B**). The best selected model in both species pairs was the secondary contact model with two categories of loci experiencing different rates of introgression (SC2*m*) and occurring in proportions *P* (for freely introgressing loci) and 1-*P* (for loci with reduced effective migration rates) across the genome. Each plot shows the observed JAFS (upper left), the JAFS obtained under the maximum-likelihood SC2*m* model (upper right), the spectrum of residuals with over (red) and under (blue) predicted SNPs (lower left), and the distribution of residuals (lower right).

Therefore, the semi-permeable species-barrier model, in which introgression occurs at variable rates across the genome since secondary contact, explains well the observed data. In the neutrally introgressing fraction of the genome (*i.e.* 55 to 63% of the genome), gene flow was found to occur in both directions at closely similar rates corresponding to 12 to 18 migrants per generation (*N*_e_*m*, Table 2). By contrast, there was a 6 to 11-fold decrease in gene flow (*m*/*m’*, Table 2) in the remaining fraction of the genome, due to reduced effective migration rates in both directions. Time parameters indicated that the duration of isolation without gene flow was 11 to 21 longer than the duration of secondary gene flow (*T*_S_/*T*_SC_, Table 2). These results support that the allopatric divergence period was long enough to enable differentiation prior to secondary contact.

## Discussion

### An improved scaffolding of the Pacific oyster genome

Our new high-density linkage map in *C. gigas* adds to a series of first-generation linkage maps (Hubert & Hedgecock, 2004, Sauvage et al., 2010, Hubert et al., 2009, Plough & Hedgecock, 2011, Guo et al., 2012, Li & Guo, 2004, Zhong et al., 2014a) and more recent second-generation maps (Hedgecock et al., 2015, Wang et al., 2016) that were produced in the Pacific oyster. Compared to the two most recent linkage maps, this map offers a slightly improved resolution, with an average marker-spacing of 0.64 cM compared to 1 cM in Hedgecock et al. (2015) and 0.8 cM in Wang et al. (2016). We paid attention to minimize the influence of the potentially widespread transmission ratio distortions observed across most of the 10 linkage groups (Plough, 2012, Plough & Hedgecock, 2011, Launey & Hedgecock, 2001, Hedgecock et al., 2015), by discarding distorted markers prior to map reconstruction. Our map, however, shows a good collinearity with the map produced by Hedgecock et al. (2015) that did not exclude distorted markers. This suggests that distortions of segregation ratios do not substantially affect the ordering of markers in *C. gigas* genetic maps, and further support that Hedgecock‘s consensus linkage map and ours provide a reasonable assessment of marker order and genetic distances across the Pacific oyster genome.

Consistent with what has been previously reported by Hedgecock et al. (2015), we found a high incidence of chimeric scaffolds (38.5% and 44.2%, respectively) in the oyster_v9 assembly (Zhang et al., 2012; http://www.oysterdb.com). Splitting scaffolds to resolve mapping conflicts to different linkage groups decreased the scaffold N50 by 45.6% (401 to 218 kb), but the rescaffolding step using BES increased the scaffold N50 again by 47% (320 kb). Due to the limited density of markers in our linkage map, this new assembly version (available at ftp://ftp.ifremer.fr/ifremer/dataref/bioinfo/) is not complete and tends to exclude especially the shortest scaffolds present in the oyster_v9 assembly (Zhang et al., 2012). Therefore, only 50.1% (280.2 Mb) of the oyster_v9 assembly could be anchored into 10 pseudo-chromosomes based on the information contained in our new linkage map. This underlines the need for an improved genome assembly in *C. gigas* using new information from higher density linkage maps and long read sequencing data. Such improvements are mandatory for all functional analyses relying on a complete genome description and a thorough annotation of gene sequences, such as genome-wide association studies (GWAS). However, for the purpose of this study, a partial chromosome-level assembly is sufficient to explore broad-scale genomic variation in genetic differentiation between the two studied sister taxa and its possible relationship with recombination rate variation.

### Recombination rate variation across the genome

The total length of our sex-averaged consensus map (965 cM) is intermediate to previous estimates that were reported from the two existing high-density linkage maps in the Pacific oyster (890-1084 cM, Hedgecock et al., 2015, respectively, Wang et al., 2016). Considering the genome-size estimate of 559 Mb (Zhang et al., 2012), this corresponds to an average recombination rate of 1.73 cM per Mb, a value within the range of average recombination rate values obtained across a wide diversity of animal species (Corbett-Detig et al., 2015, Stapley et al., 2017).

The chromosome-averaged recombination rate shows a 2.5 fold variation among linkage groups and is negatively correlated to chromosome length. Such a pattern has already been observed in several species including yeast (Kaback et al., 1992), human (Lander et al., 2001), stickleback (Roesti et al., 2013) and daphnia (Dukić et al., 2016), and has been commonly attributed to positive crossover interference. This phenomenon happens during meiosis, when the formation of an initial recombination event reduces the probability that an additional recombination event occurs nearby on the chromosome. If the mechanism of interference is still elusive, it is often assumed that a fixed number of crossovers happens per chromosome (between 1 and 2), making large chromosomes experiencing less recombination events per nucleotide position than smaller ones. Our results thus provide new empirical support for the existence of crossover interference in *C. gigas*.

We also revealed large-scale (i.e. megabase-scale) variation in local recombination rate within chromosomes. The general pattern we observed was a reduction of crossover rate in chromosome centers relative to peripheries. Similar patterns are widespread in animals, plants and fungi (reviewed in Berner & Roesti, 2017), and can be attributed to different but not mutually exclusive mechanisms. Firstly, a reduction in recombination near the centromere of metacentric chromosomes (Nachman, 2002) can be hypothesized, possibly due to the presence of highly condensed heterochromatin. Secondly, crossover interference, which tends to produce evenly and widely spaced crossovers when multiple crossovers occur on the same chromosome, may increase the chance that multiple recombination events affect chromosomal extremities. Thirdly, heterochiasmy (i.e. different recombination rates and landscapes between sexes) could also explain these patterns (Stapley et al., 2017), especially if one sex tends to recombine preferentially in chromosomal extremities. In the Pacific oyster, the comparison of male and female maps did not support the hypothesis of heterochiasmy (Hedgecock et al., 2015), so the observed recombination rate variation within chromosomes could be rather explained by centromere location and/or crossover interference.

The recombinational environment is known to affect the impact of natural selection on linked neutral diversity across a wide range of species, in which low-recombining regions generally display reduced levels of nucleotide diversity (Corbett-Detig et al., 2015). As expected under the joint effect of background selection (Charlesworth et al., 1993) and hitchhiking (Maynard Smith & Haigh, 1974) on linked neutral diversity, we found a positive relationship between the local recombination rate and nucleotide diversity across the Pacific oyster genome. However, the magnitude of this effect was rather small compared to what could be expected from comparative estimates in other invertebrate species supposed to have large population sizes. (Corbett-Detig et al., 2015) argued that the effect of natural selection on the reduction of linked neutral variation was stronger in species with large census sizes, but a reanalysis by (Coop, 2016) showed that this conclusion was not supported by the data. In addition, other invertebrate species studied so far tend to have smaller genomes than the oyster genome. Oysters are usually perceived as species with large effective population sizes and this is corroborated by a medium to high nucleotide diversity (Sauvage et al., 2007, Zhang et al., 2012). However, several empirical evidences from molecular evolution and population genetics studies have also shown that oysters have among the highest segregating loads of deleterious mutations observed in marine invertebrates (Sauvage et al., 2007, Plough, 2016). This may be due to a high variance in reproductive success (sweepstake effect) and population size fluctuations (Boudry, et al. 2002; Harrang, et al. 2013; Hedgecock 1994; Hedgecock and Pudovkin 2011; Plough 2016). We are lacking theoretical predictions on the effect of linked selection in a species with skewed offspring distribution, but the effect is likely more genome-wide during favorable sweepstake events than it is in the standard Wright-Fisher model.

### Genome-wide diversity patterns within and between Pacific oyster species

The core objectives of this study were to (*i*) determine the extent and variation in the level of differentiation within and between *C. angulata* and *C. gigas* across the genome, (*ii*) test the existence of differences in the genomic landscape of differentiation between native and introduced species pairs, *(iii)* infer the history of gene flow during divergence, and finally (*iv*) relate interspecies divergence to underlying evolutionary processes including demography, selection and genomic constraints.

Our results reveal that the European introduction of *C. angulata* in the 16^th^ century, and more recently of *C. gigas* in the 70’s, did not lead to a reduction in genetic diversity for both species compared to source populations in their native range. This observation is rather common in the literature of marine invasions, and has been attributed to the combined effects of multiple introductions and high propagule pressure (Viard et al., 2016). Another possibility is that the initiation of the demographic wave of invasion can simply not occur without a sufficiently large number of individuals in species with a strong Allele effect like sessile broadcast spawners. In other words a successful marine introduction cannot happen without a high number of funders. We found a relatively low background level of differentiation between *C. gigas* and *C. angulata* (i.e. genome-wide averaged *F*_ST_ = 0.045), which is consistent with previous estimates based on microsatellite loci (Huvet et al., 2004, Huvet et al., 2000). However, differentiation was highly heterogeneous across the genome in both Asian and European populations, with peaks of elevated *F*_ST_ values being found in most chromosomes, sometimes even reaching differential allelic fixation in Asia. These ‘genomic islands of differentiation’ are commonly observed in genome scans for divergence between closely related species pairs (Wolf & Ellegren, 2017), and multiple mechanisms have been proposed to explain their formation (Cruickshank & Hahn, 2014, Burri et al., 2015, Ravinet et al., 2017, Yeaman et al., 2016). They broadly fall in two categories: (*i*) mechanisms of differential gene flow involving the existence of reproductive isolation and/or local adaptation loci acting as genetic barriers to interspecific gene flow (Barton & Bengtsson, 1986, Feder & Nosil, 2010), and (*ii*) heterogeneous divergence mechanisms due to variation in the rate of lineage sorting across the genome (Burri et al., 2015, Cruickshank & Hahn, 2014, Burri, 2017).

In order to test whether specific adaptations to different habitats participate to genetic barriers, we hypothesized that the different ecological conditions encountered by oysters in Europe would either result in relaxed divergent selection pressures on genes involved in local adaptation in Asia, or in new selective constraints targeting different subsets of genes. In both cases, we expect that the new environmental conditions in Europe would promote a remodeling of the genomic landscape of species divergence and hence reduce the extent of parallelism in genetic differentiation between species-pairs from native and introduced populations. Contrary to that prediction, we found a remarkably high degree of divergence parallelism indicating that the genetic architecture of reproductive isolation between *C. angulata* and *C. gigas* has not been reinforced, nor significantly weakened following the co-introduction of the two species in Europe. Nevertheless, a slightly lower interspecific differentiation was found in Europe. This reduction was not driven by the most differentiated loci (Fig. 3B), and therefore rather indicated increased gene flow across the whole genome than relaxed divergent selection on barrier loci in the novel habitat. Moreover, the result of the PCA suggests that more pronounced gene flow is ongoing from *C. gigas* to *C. angulata* than in the opposite direction in Europe, as revealed by the more pronounced shift of *C. angulata* samples from Portugal towards the central part of the first PCA axis compared to *C. gigas* samples from Brittany (Fig. 3A). This asymmetrical introgression pattern seems consistent with a demographic imbalance due to a higher abundance of *C. gigas* in European farms. Overall, these results suggest that the particular demographic conditions imposed by aquaculture in southern Europe have facilitated opportunities for genetic interactions between species, without modifying the nature of the species boundary.

The finding of unaltered genomic differentiation patterns in the introduced species-pair despite increased gene flow suggests either that the contact is too recent to observe significant effects, or that historical and architectural genomic features have played major role in shaping the genomic landscape of species divergence between Pacific cupped oysters. Moreover, heterogeneous genome divergence is predicted by the semipermeable species boundary model, in which neutral introgression is only permitted in genomic regions unlinked to reproductive isolation barriers (Harrison & Larson, 2014, Harrison & Larson, 2016). Under this model, elevated *F*_ST_ values above the background level indicate the location of genomic regions that are resistant to introgression due to the presence of reproductive isolation loci (i.e. speciation genes). The existence and the possible origin of a semi-permeable barrier to gene flow between *C. gigas* and *C. angulata* was addressed through the comparison of various theoretical models offering simplified representations of contrasted evolutionary scenarios. Our results indicated that contemporary differentiation patterns likely result from a long period of allopatric divergence followed by a recent secondary contact with different rates of introgression among loci. This model consistently outperformed alternative scenarios in both native and introduced ranges, predicting observed differentiation patterns with very small residual errors (Figure 6). Inferred model parameters capturing semi-permeability indicated that the fraction of the genome experiencing reduced introgression rates amounts to 37-45 %, consistent with the view that the species barrier is still largely permeable to gene flow. Similar estimates have been obtained in other marine species-pairs analyzed with the same approach (Tine et al., 2014, Le Moan et al., 2016, Rougemont et al., 2017). In all these cases post-glacial secondary contacts between allopatrically diverged lineages have been inferred. The northwestern Pacific region is known to harbor multiple cases of marine species subdivided into divergent lineages that are supposed to have originated in separate glacial refugia, most likely corresponding to the current seas of Japan and China (reviewed in Ni et al., 2014). Our study provides the first genome-wide view of the evolutionary consequences of past sea-level regression in this region, and adds to previous studies suggesting their important role in initiating speciation in northwestern Pacific taxa (e.g. Shen et al., 2011).

In addition to providing evidence for partial reproductive isolation between *C. gigas* and *C. angulata*, our results also bring new empirical support for the role of recombination rate variation in shaping the genomic landscape of species divergence. A recent study in the European sea bass has shown that low-recombining regions experiencing accelerated rates of lineage sorting during allopatric phases are preferentially involved in the barrier to gene flow during secondary contact (Duranton et al., 2017). The negative relationship between differentiation and recombination in Pacific cupped oysters is consistent with the idea that foreign alleles are more efficiently removed by selection after introgression when recombination is locally reduced (Schumer, et al. 2017). This interpretation also suggests that the sites under selection are widespread across the genome, although not individually under strong selection (Aeschbacher et al., 2017). Evidence for this type of genetic architecture has recently been evidenced from hybridization studies in a number of animal and plant species (Simon et al., 2017).

## Conclusion

Our results shed new light on the existence of reproductive isolation barriers between the two cupped oysters *C. angulata* and *C. gigas.* By providing empirical evidence for heterogeneous divergence patterns attributable to reduced introgression in low-recombining regions since secondary contact, we show that these semi-species are still evolving in the so-called “speciation grey zone” (Roux et al., 2016). Moreover, the finding of strong divergence parallelism between species-pairs from native and introduced areas suggests that the genomic architecture of reproductive isolation is not primarily determined by ecological divergence driven by local adaptations in the native range. Our study thus implies the existence of intrinsic genetic differences between the two species, which will be the focus of future investigations based on experimental crosses.

## Acknowledgements

This work was supported by Interreg SUDOE “Aquagenet” (SOE2/P1/E287) and ANR “GAMETOGENES” (ANR-08-GENM041) projects, as well as from GIS-IBiSA and Ifremer. We would like to thank Brieg Pontreau for his help in microsatellite genotyping, the hatchery team of Ifremer La Tremblade for its help in the production of the biological material, and Hélène Holota from Inserm UMR 1090 TAGC for her kind assistance during the sonication of RAD libraries. We are also grateful to Patrick Gaffney for providing BAC clones and Pascal Favrel for managing BAC-end sequencing with the Genoscope. Golden Gate SNP genotyping was performed at the Genome Transcriptome Facility of Bordeaux (grants from Conseil Régional d’Aquitaine n°20030304002FA and 20040305003FA, from the European Union FEDER n°2003227 and from the Investissement d’Avenir ANR-10-EQPX-16-01). Sequencing was performed by the Genoscope, Integragen and the CNRS plateform Maladies Métaboliques et Infectieuses.

